# Marching with ants to a new nest

**DOI:** 10.1101/2023.01.27.526000

**Authors:** Sumana Annagiri, Eshika Halder

**Affiliations:** Behaviour and Ecology Lab, Department of Biological Sciences, Indian Institute of Science Education and Research Kolkata, Mohanpur, West Bengal 741246, India

**Keywords:** Ponerinae, *Diacamma indicum*, Colony composition, Gamergate, Tandem running, Brood theft, Path optimization

## Abstract

In this review, we journey with *Diacamma indicum* a Ponerine ant over the last decade as they relocate to new nests and discover the challenges they face along the way and how they solve them. Colony relocation is a goal oriented dynamic task that involves all the colony members and impacts the colonies’ fitness. After explaining how I initiated this journey, we examine colony composition of this species by analysing data from 1200 colonies collected over the last 13 years. On average colonies are constituted with 89.35 adult females, 0.29 males and 56.6 brood items of different development stages and these were significantly impacted by seasonality with Pre monsoon having the highest numbers. After explaining how colonies are collected and maintained in the lab, we explore the architectural components of the subterrain nests built by this species in the natural habitat. Colonies live in relatively simple single chambered nest that does not change significantly across seasons and consists of an entrance tunnel and a secondary runoff tunnel. All members of the colony are recruited to the new nest through tandem running and this species shows the highest documented tandem running speeds at 4.35 body lengths per second and a path efficiency of 83.95% with only 2.4% of tandem runs being unsuccessful in the natural habitat. Even in lab conditions, when colonies are given defined paths of different lengths, colonies showed significant preference to travel through short paths, highlighting their ability to optimizes their path even in the absence of chemical trails. A combination of experiments in the natural habitat and controlled experiments in the lab which are anchored in the *umwelt* of the organism has enabled us to understand how *D. indicum* functions and reveals the selection forces that are operating on the organization and performances of relocation. Our journey has brought to light several answers but has opened several more avenues for exploration branching out in different directions. With time and dedicated minds, we hope to continue on this route to marvel at the achievements of these superorganisms.

## Initiation

It was late morning and the sun was getting strong enough to dismiss the cold blanket that had enveloped the grounds. A pair of black ants were marching right past my foot and after about a meter or so they entered into the drain hole of a potted palm plant that was placed in our front veranda. It was not a trail of ants just the two of them. After a couple of minutes, an ant emerged from the same hole and walked out, while I was putting my shoe on. But before I got ready to leave the house, I saw another pair of ants marching into the same destination. It was really intriguing, where was the trail, would it be the same ant going back and forth, what was happening? I had to go out as there was a meeting to attend, but marching ants stayed in my mind that day and have been on my mind for the past 13 years now.

Having joined the workforce after a four-year break, I was teaching Animal behaviour to the first batch of students at our Institute and looking for questions to initiate my research. Insects particularly those that can live in groups were of primary interest to me, but I was open to exploring and finding a new niche for my lab. This is not as easy as it may sound as the path is filled with uncertainties and the stakes are high. Having moved back from USA into a beautiful but remote corner of Bengal we faced several challenges. Surrounded by a new culture, new language, several well-meaning relatives, a little daughter who was starting to explore the world and an Institute that was initiating its journey. Arguably this shift from USA to Kolkata caused a significantly higher degree of perturbance than my earlier move from India to Boston.

Even though I had never worked with ants, I had seen other people working with ants when I was doing my Ph.D., I took a shovel, a pair of gloves and a collection flask and thought of collecting the ant colony. While I was working on it, our gardener stopped by and said that I was doing it wrong – the digging part. So, I explained that I was not doing gardening but looking for the nest, he certainly was not impressed either with my Bengali or my technique of digging but he hung around together with lots of advice. Once he saw the ants coming out of its nest he said “oh these ants, near my house we have this and much bigger ones!” As these ants are very much a part of the everyday life of the people around here, they are full of stories about them and what they think the ants are doing. This naturally led to several questions – does scale matter, bigger the better? Of course not, a naturalist would say, each organism is fascinating in its own right. Does the location that it occurs matter? Our backyards are as fascinating as the remote corners of this landmass, as a world anew is waiting to be looked into in both places. So, I went ahead, digging several spades of mud I got colonies back to the lab. Lab being one table next to a centrifuge in the centre of an instrument room in the newly renovated building. Once I could get ants to march in pairs within the lab, I had a hunch that I had a potentially interesting question that I can pursue. Preliminary investigations led me to believe that I had an ant belonging to genus *Diacamma* and with some help from a taxonomist we identified that it belonged to the species *indicum*. This 1 cm black ant, with metallic hues on its body, a head with prominent eyes, 12 segments in its antennae and no ocelli, a prominent petiole with two spines (Figure 1A). Searching literature in the initial days showed that there was a grand total of 3 research papers on this species, so I decided to go ahead with my explorations and my own march with these ants.

**Figure 1.**
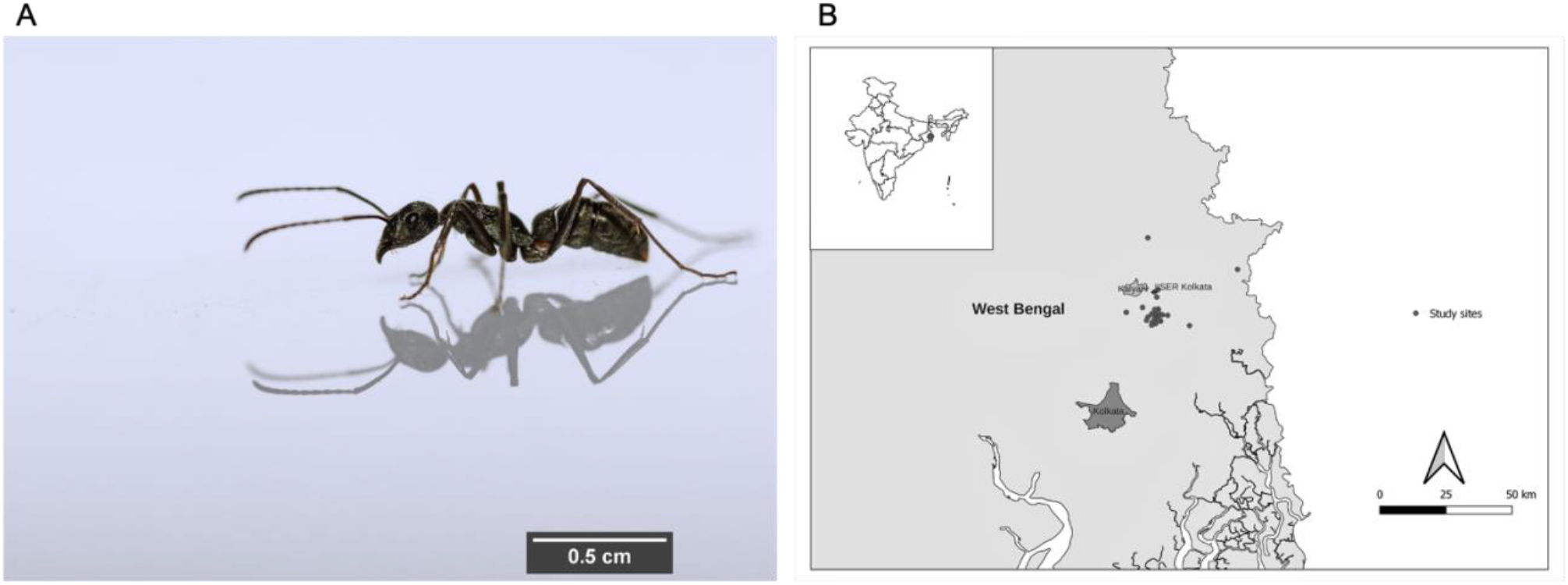
A)Female *Diacamma indicum* taking a stroll. (Picture courtesy – Subhashis Halder) B) The location of the study area is depicted on the map of India.

In India, a total of 111 species belonging to the Ponerine taxa have been reported. Out of that 12 species of *Diacamma* are found in India and 6 species of *Diacamma* are reported in West Bengal (Bharti et al., 2016). Several studies have been conducted on Ponerine ants which have mostly dealt with foraging and particularly how they hunt for termites (Dejean, 2011; Fowler, 1980; Pie, 2004). *Diacamma* is particularly interesting because of the unique method by which they achieve reproductive monopoly in their colonies (Annagiri, 2021; Peeters & Higashi, 1989). But this will be a story to tell on another occasion.

## Study area

All the *Diacamma indicum* colonies were collected from the districts of Nadia (23.4710° N, 88.5565° E) and North 24 Paraganas (22.7271° N, 88.7298° E), West Bengal. This region is situated on the eastern side of West Bengal, India (Figure 1B). It is approximately 50 km away from the capital of the state, Kolkata. The district is surrounded by the Bhagirathi River in the west and the Ichhamati river in the north. The entire district lies on the Gangetic plain consisting of alluvial soil. This tropical region receives an annual rainfall of 200 cm during monsoon (June to September). The temperature ranges from 10°C to 41°C and the humidity range from 44% to 87%. It is one of the agriculturally developed areas with a high diversity of flora and fauna. However, in the last two decades, it is experiencing urbanization, industrialization and a lot of fishing activities (Biswas et al., 2015)

## Colony composition

From about 1200 colonies that we have collected to date, we characterized all the data for the first time and presented some natural history facts regarding the colony composition of this species here. Colony size as measured by the number of adult females in the colony on average is 85.35 (standard deviation is 38.79). Is there a variation in colony size across the different months in the year? In this tropical region, where temperatures do not fluctuate very much one of the major abiotic factors that impact the habitat is the monsoons. Thus, it was meaningful to classify the year into three different seasons based on monsoons. Four months that is July to October is considered as monsoon season, while November to February is considered as Post monsoon and March to June as Pre monsoon months. (Kolay & Annagiri, 2015). The number of colonies collected during pre-monsoon, monsoon and post monsoon was N_1_ = 443, N_2_ = 441, N_3_ = 309 respectively. Colony size changes significantly over seasons with the pre-monsoon months having the highest number of adults (92.75 ± 38.47) as compared to monsoon (90.9 ± 39.96) and post monsoon (82.26 ± 36.75) (Kruskal-Wallis test: N_1_ = 443, N_2_ = 441, N_3_ = 309, p < 0.05) (Figure 2A). By ant standards, these numbers were certainly at the lower end of the spectrum as other species of ants can have millions of individuals (Dornhaus et al., 2012). Colonies almost always had a single individual with gemma and this individual is known to be the sole reproductive of the colony and is termed as the gamergate (Peeters & Higashi, 1989). Of the colonies collected across different seasons, it was seen that significantly higher number of colonies did not have its gamergate during the post monsoon season (Chi-squared: 13.11, p < 0.05). The number of colonies where the gamergate was missing in pre monsoon, monsoon and post monsoon were 31, 29 and 43 respectively. Interesting males were present in colonies throughout the different seasons in low numbers and there was no significant difference in their occurrences across seasons which would indicate that this species does not have a well-defined mating season (Kruskal-wallis test: N_1_ = 443, N_2_ = 441, N_3_ = 309, Kruskal-Wallis chi-squared = 1.9, p = 0.38) (Figure 2B). The number of broods that is eggs, larvae and pupae were significantly different across seasons (Kruskal-Wallis test: N_1_ = 443, N_2_ = 441, N_3_ = 309, Kruskal-Wallis chi-squared = 141.88, p < 0.05) (Figure 2C). The highest and lowest number of brood were found in pre monsoon (61.80 ± 47.89) and post monsoon (35.06 ± 31.39) respectively. The number of brood in the colony was strongly correlated to the number of adult females in the colony across seasons (r = 0.5, p < 0.05, GLM: McFadden r^2^ = 0.24, y = 0.008x + 3.19) (Figure 3A). The relative proportion of the three different development stages that is egg, larva and pupa were 1:0.67:1.48 in pre monsoon, 1:0.63:1.42 in monsoon, and 1:0.59:2.03 post monsoon respectively (Figure 3B). The number of pupa was higher during monsoon (21.79 ± 16.54) as compare to pre monsoon (19.61 ± 18.32) and post monsoon (9.66 ± 13.72) (Kruskal-Wallis test: N_1_ = 443, N_2_ = 441, N_3_ = 309, Kruskal-Wallis chi-squared = 161.71, p < 0.05). The number of larva was higher during monsoon (13.67 ± 10.89) as compare to pre monsoon (13.13 ± 11.69) and post monsoon (5.73 ± 7.29) (Kruskal-Wallis test: N_1_ = 443, N_2_ = 441, N_3_ = 309, Kruskal-Wallis chi-squared = 153.11, p < 0.05). The number of eggs was also higher during monsoon (31.67 ± 20.85) as compare to pre monsoon (29.06 ± 25.31) and post monsoon (19.66 ± 18.12). (Kruskal-Wallis test: N_1_ = 443, N_2_ = 441, N_3_ = 309, Kruskal-Wallis chi-squared = 62.66, p < 0.05)

**Figure 2:**
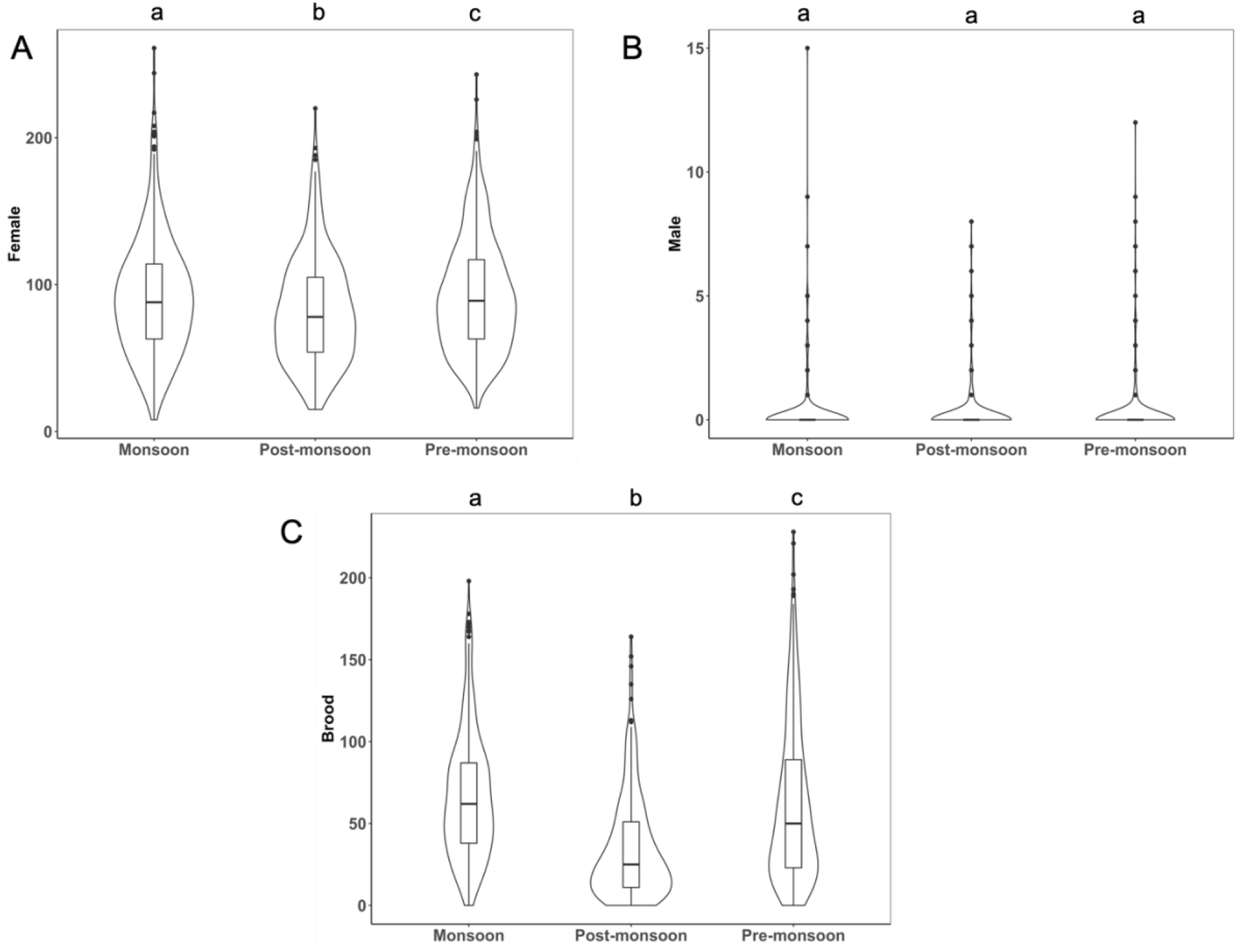
Number of adult females, males and brood across seasons in colonies: This violin plot depicts A) The number of females present in a colony across three seasons (N_1_ = 443, N_2_ = 441, N_3_ = 309, p < 0.05). B) The number of males present in colonies across three seasons (N_1_ = 443, N_2_ = 441, N_3_ = 309, p = 0.38). C) The number of broods present across three seasons (N_1_ = 443, N_2_ = 441, N_3_ = 309, p < 0.05). The bold black line inside the box represents the median value and the box represents the interquartile region. The area under the curve represents the density distribution of the data. Comparison has been done using Kruskal-Wallis test and post hoc Dunn test with holm correction

**Figure 3.**
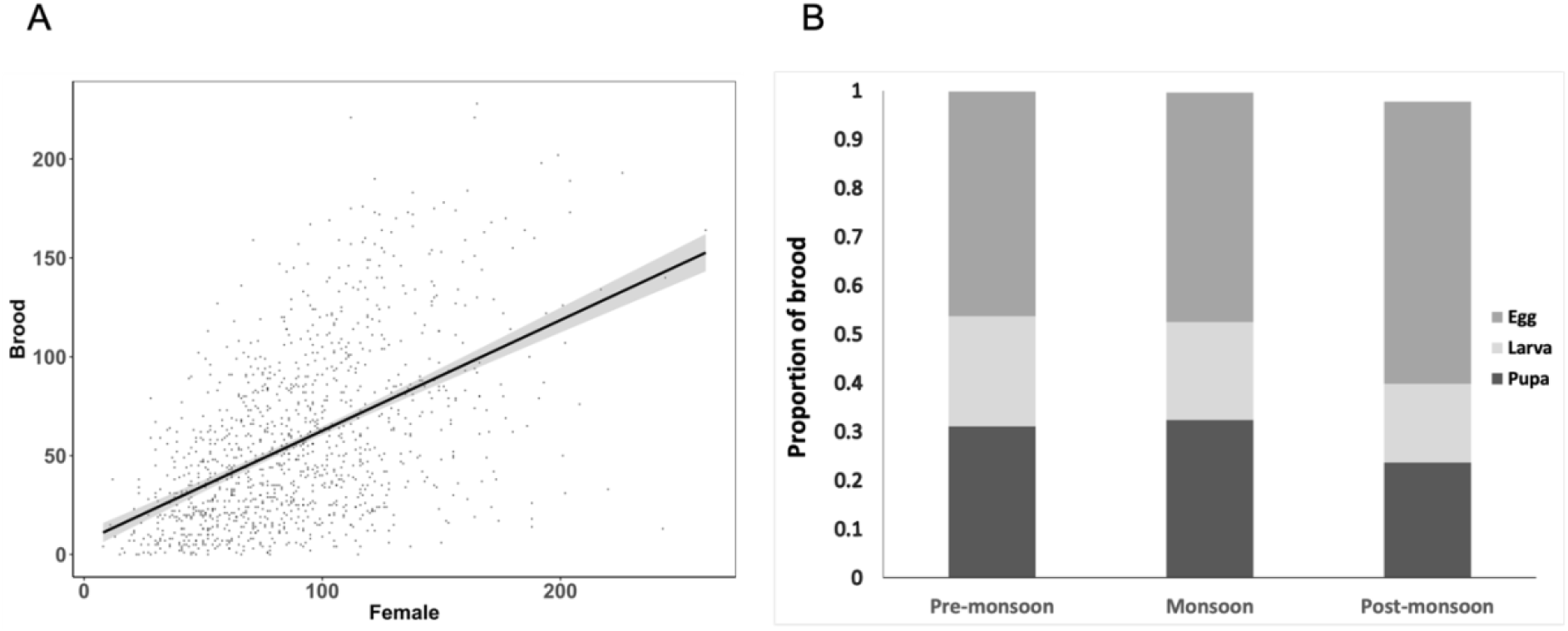
A. Brood in the colony: Relationship between the number of brood and adult females in a colony. A significant correlation between the number of females (x axis) and number of brood in the colony (y axis) was detected (N =1193 colonies, Spearman correlation, r = 0.5, p < 0.05, GLM, McFadden r^2^ = 0.24, y = 0.008x + 3.19 is plotted as the black line). The grey region surrounding this black line represents 95% confidence interval on the fitted values. B) The proportion of brood in the colony: The stacked histogram depicts the changes in the proportion of brood across three seasons. Pre monsoon (N_1_ = 443), monsoon (N_2_ = 441) and post monsoon (N_3_ = 309).

Overall, colony composition was impacted by seasons and the presence of gamergate. The presence of gamergate has a significant positive impact on the number of females present in the colony (GLM, est = 0.3, p < 0.05). Pre monsoon (GLM, est = 0.02, p < 0.05) has a positive impact on the number of females. However, post monsoon (GLM, est = −0.07, p < 0.05) has a significant negative impact on the number of females (Fig 2A). The number of brood present in the colonies was significantly positively affected by the presence of gamergate (GLM, est = 0.4, p < 0.05), negatively affected by seasonal variation (pre monsoon, GLM, est = −0.08, p < 0.05, post monsoon, GLM, est = −0.5, p < 0.05) (Fig 2C), and positively by the number of females (GLM, est = 0.007, p < 0.05) (Fig 3A). The number of males was comparable across season (pre monsoon, GLM, est = 0.11, p = 0.33, post monsoon, GLM, est = 0.13, p =0.32) (Fig 2B). The details of the generalized linear model used has been provided as supplementary material.

## Colony collection by nest flooding method

After collecting several colonies by removing the surrounding soil in the traditional manner, we had a preliminary idea of the typical nest for this species. Nests had a single entrance and a single nest chamber and were situated within 30 cm depth from the surface. Equipped with this information my first Ph.D. student decided to try a new method to collect colonies (Kaur, 2014). A plastic box was connected to plastic tube with a diameter that is similar to the nest entrance. This plastic tube was inserted into the nest entrance to the depth that it would go without causing damage inside the nest. Any gaps at the entrance of the nest and the collection box were plugged with cotton so that ants could not escape the setup (Figure 5A). Once this was in place the nest entrance received water in a slow but steady stream so that it entered the entrance tunnel and the chambers. This water stress caused ants to evaluate the flooded nest chamber and walk into the tube which was dry on the inside to the collection flask. Several of these ants held brood in their mandibles as they escaped. Some ants were seen making return trips to the chamber and perhaps bring additional brood which was deposited into the collection flask. Once the rate at which ants walked into the tube decreased the rate at which the water was poured into the nest was increased. After a couple of minutes of this flooding, no further ants were seen entering the tube from the nest and at this stage the plastic tube is detached from nest entrance and plugged with cotton. In the interim, any returning forgers were collected and placed inside a separate collection flask to be reunited with the colony inside the lab. In the next step, water was poured into the nest entrance directly and we waited for any ants to emerge from the entrance, if no ant emerged the whole nest is flooded until water rises to the entrance to ensure that the chamber is empty. This method allowed us to collect colonies with greater ease and mimic a natural event that can possibly occur during monsoons, when rain enters into their nest and cause flooding. This method has been effective across the years and can be recommended for not only *D. indicum*, but other species of ants that have a small number of known nest entrance and a single chamber which is not very deep. Species *like Odontomachus brunneus* (Cerquera & Tschinkel, 2010), *Pheidole oxyops* (Forti et al., 2007), *Trachymyrmex septentrionalis* (Tschinkel & Bhatkar, 1974), can be considered as candidates for testing this method of colony collection as they have a subterranean nest structure that is similar to *D. indicum*.

**Figure 5.**
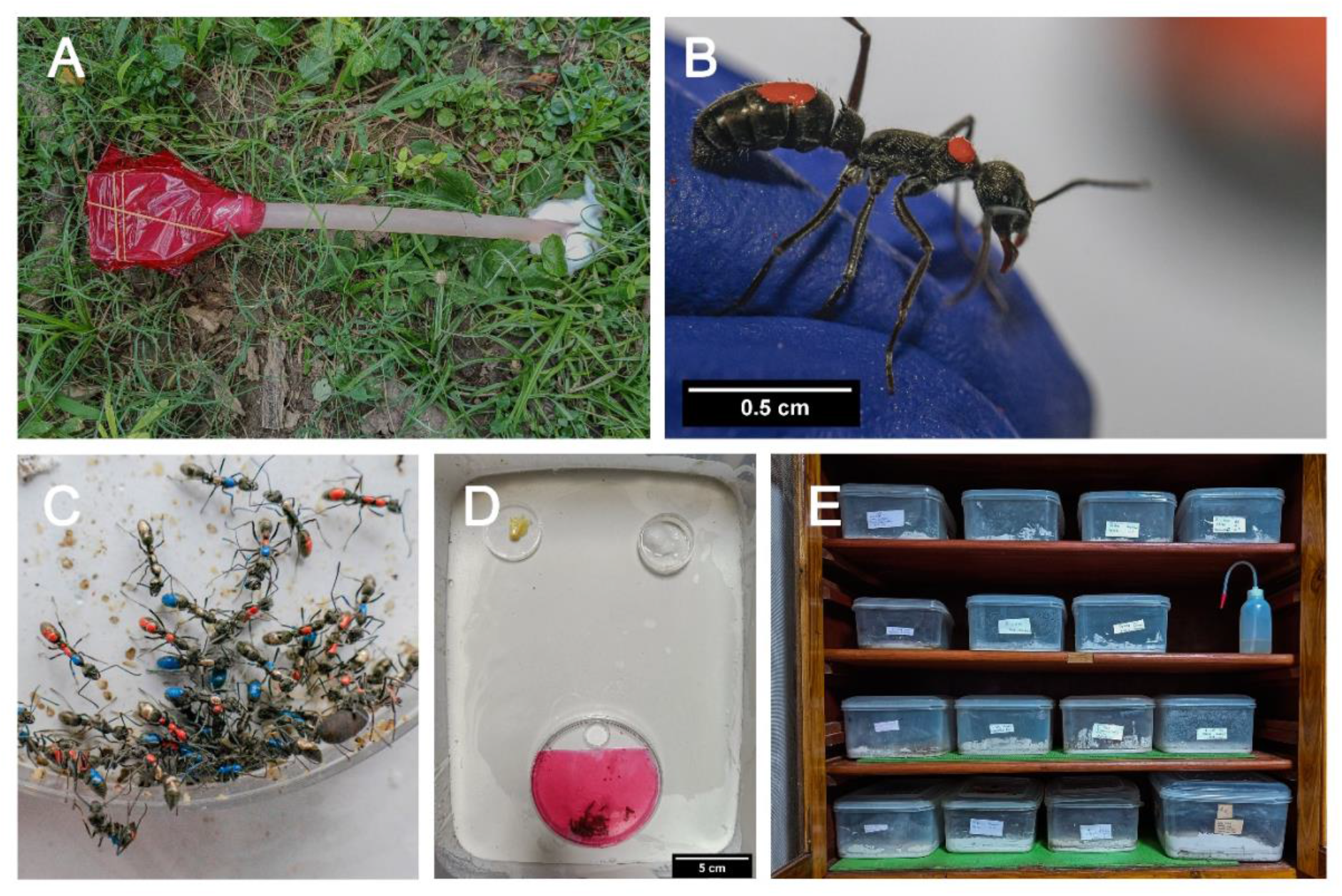
Pictures depicts the process by which ant colonies are collected and maintained in the lab. (A) shows colony collection from their natural habitat using the water flooding method. (B) Each ant within the colony is marked with enamel paint on different parts of their body for individual identification by the experimenters. Marking this particular ant inside as red blank red or R-R in short. (C) Colonies are housed in these nest boxes (28.5cm X 21.5cm X 12cm) inside the artificial nest consisting of plastic petri plates. (D) Colonies are typically provided *ad libitum* food and water in small petri plates inside the nest box as seen in the picture. (E). Colonies holding such nest boxes are maintained inside the ventilated wooden almirah. (Pictures courtesy – Subhashis Halder).

## Living in the Lab

Colonies are housed in nest boxes (28.5cm X 21.5cm X 12cm). These plastic boxes have a plaster of Paris base and ant colonies live inside an artificial nest consisting of plastic petri plates. The bottom plate 9 cm is coated with plaster of Paris and the top plate which acts as a roof has a circular entrance (90mm diameter) cut into it. These colonies are typically provided *ad libitum* food (Bhatkar & Whitcomb, 1970)_and water in small petri plates inside the nest box (Figure 5D). Further, these nest box’s walls are coated with Vaseline (Hindustan Unilever Limited, India) in order to prevent ants from escaping the setup. Following that, we count every individual and the number of broods present in the colonies and enter the same into a lab register (Karunakaran & Annagiri, 2018; Sumana & Sona, 2013). Each ant within the colony is marked with enamel paint (Testors, Rockford, IL, USA) on different parts of their body for individual identification by the experimenters. Remember, all members of the colony look uniformly black with metallic hues to our eyes and there is no way we can distinguish between individuals. In order to overcome this issue, we mark the ants. While the ant is gently held with two fingers a small dot of the required colour is placed on the body of the ant marking them (Figure 5C, which shows an ant being marked as red blank red or R-R in short). Typically, we use three areas on the body of the ant to make these marks, hence we say blank or – is one of the areas that does not have a colour. This way we can other ants marked as RR-, RRR, R— and so forth. When we use a combination of colours you can image that we will indeed have “colourful” colonies with unique ants (Figure 5C). Following this step and at least one day time for them to acclimatize to the lab conditions colonies are considered ready for experiments. Colonies holding such nest boxes are maintained inside ventilated wooden almirah (Figure 5E).

## Tandem running

It turned out that the pair of ants that were walking with almost constant contact with each other had been documented before, as early as 1896 by Gottfrid Adlerz. Several researchers had studied it in detail in other species such as *Temnothorax albipennis, Camponotus sp, Harpagoxenus sp* (Franks & Richardson, 2006; Schultheiss et al., 2015; Stuart & Alloway, 1985). This recruitment method is used in the context of relocation, foraging and conducting slave raids on neighbouring colonies (Franklin, 2014). Hardly anything was known about *Diacamma indicum* and nothing about its tandem running behaviour had been documented. The ant in front, termed leader recruits other nestmates termed follower via tandem running to the desired destination. After exploring a little more, I found out that what I had first seen was a colony relocating from their old nest into a new nest via tandem running (Figure 6). After doing a series of experiments in the lab, we discovered is that on average 17.9% of the colony become tandem leaders and they perform at least one tandem run each (Kaur et al., 2012). However, typically there are several leaders who perform just a single or couple of tandem runs while a small percentage perform the vast majority of tandem runs, with the leader who has done most of the tandem runs (termed Maximum tandem leader) doing almost quarter of the job (24.2% ± 10.5% tandem runs) within a single relocation. Yes, that is indeed a lot of back-and-forth trips, now we wanted to examine what is the distance over which they relocate and found out that on average they moved 1.4 m to their new nest and the range was 61 cm to 678 cm in the natural habitat (Kaur et al., 2012). A simple back of the envelop calculation shows that the Max tandem leader would thus have walked for about 59 m during the relocation, an incredible feat for a 1 cm ant by any standard.

**Figure 6.**
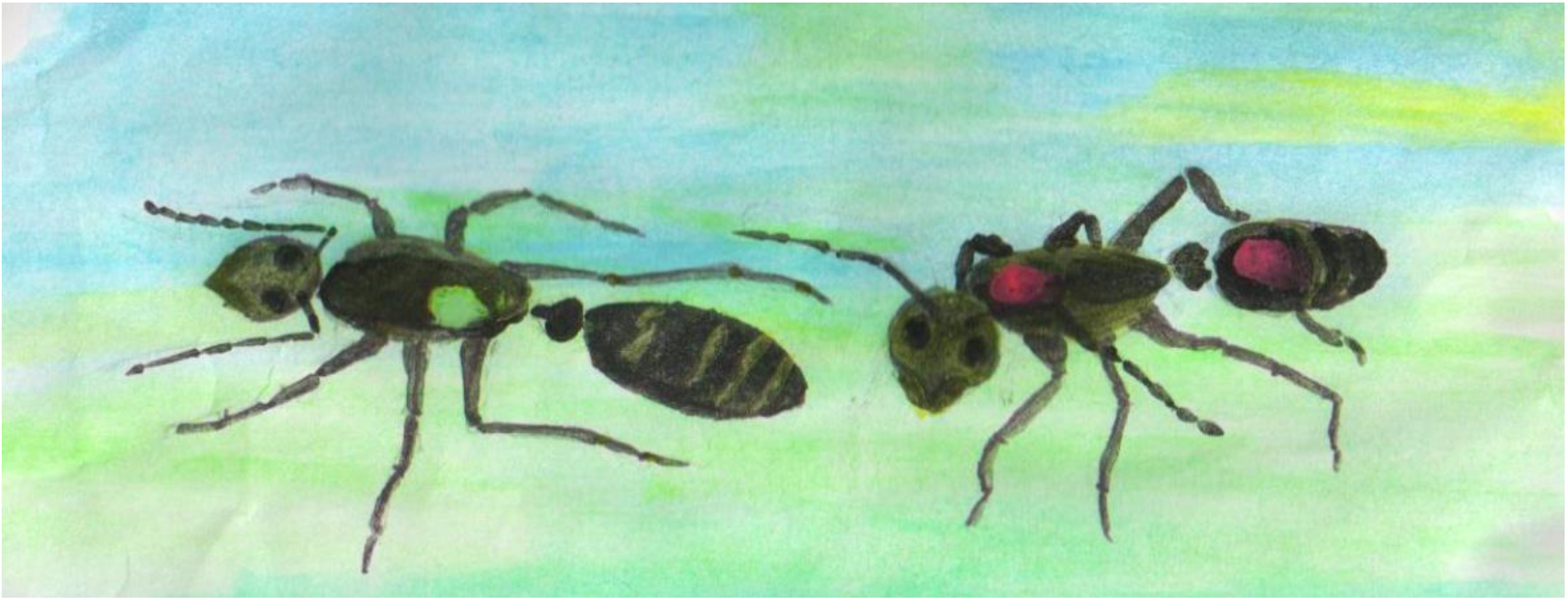
Water colour painting of tandem running *Diacamma indicum* with Green – being the leader and Red – Red being her follower. Artwork – Sumana Annagiri.

The next question of course was how fast do they go? Examining this within the lab on an artificial surface would not be relevant but it would be easier to perform. Thus, we started with that and in the next step we examined their progress in the natural habitat by following 1000 transports as they covered heterogeneous terrain to find that tandem running was indeed slower than normal walking (Anoop et al., 2021). While individual ants walked at the speed of 6.60 bl/sec (Kaur et al., 2017), tandem pairs walked at the speed of 3.81cm/sec and if the follower was holding a brood item like pupa in their mandibles while they followed a leader it was even more slow 2.74 cm/sec on average. However, followers bringing brood along with them (termed couple adult-brood transport) was more efficient because this would remove the necessity for the leader to make an additional trip to bring the brood to the new nest (Anoop et al., 2021, p. 2021; Kaur & Sumana, 2014). Further it was amazing to see the cohesion between the leader and follower. If this cohesion is weak as they walk across different terrains and vegetation over long distances, one would expect large number of interruptions as follower loses contact with the leaders. Unless the leader and follower find each other, it will result in an unsuccessful tandem run as the leader does not end up transferring her follower to the new nest. This would be a disaster because losing nestmates means losing workforce and hence fitness in the long run. As the sole reproductive of the colony – gamergate is also tandem run like to any other worker, losing her would be particularly disadvantageous to the colony. We found that only 2.4% of tandem runs were unsuccessful, in other words tandem leader lost its follower and returned to the old or new nest by herself (Anoop et al., 2021).

## Challenges during relocation

To date, I have not found a single human who has classified their experience of moving from one house to another as pleasant and something that they look forward to doing. Relocation involves a lot of uncertainties, risk and cost in terms of labour. In addition to the issues relating to scouting for a potential new dwelling, deciding on a particular option out of the several available options, settling down at a new location also needs to be taken into account. Some ant species are known to reside within the same nest for decades but others relocate more often. Physical damage to their nest is one of the major causes of relocation (Visscher, 2007; McGlynn, 2012). The organization of relocation and maintaining cohesion of colonies in these dynamic conditions in which day to day activities are disrupted also have to be considered. Then, relocating colonies would have to consider the neighbours, would there be antagonistic interactions with them? Unlike foraging where mostly experienced individuals who are aware of the landscape surrounding their nest participate, relocation involves all colony mates, including individuals who do not typically come out the nest and hence are not aware of the landscape and thus more likely to get lost making the whole process more complex. Through our studies, we investigated these issues in *D. indicum* using a series of experiments both in the natural habitat as well as in controlled conditions within the lab. In the following section, we will touch upon some of these questions and the answers we have found in a brief manner. This is only a glimpse into what we have been doing in the ant lab. Together with my students, we have just managed to open the door into the understanding of this species over the last decade but the mansion await…

### Deciding on the new nest

What are the architectural features that these ants look for in their new dwelling? Does the distance over which they have to relocate count? By conducting carefully designed choice experiments and their respective control experiments, we could answer these questions. As it turns out, in order to answer these questions initial explorations of the natural conditions of the ants dwelling were required. We first have to answer what forms a normal nest for this species. We can provide a variety of manmade nest choices but they would only be an artifact of our imagination as they lack an anchor in the realistic *umwelt* of this species. This was done by making Aluminium casts of naturally occurring nests throughout the year to find that these species live in simple subterranean nests. Monodomous, single chamber nest which is 15.9 cm deep on average. It has a secondary tunnel going deeper into the soil (Figure 7). After quantifying various features and analysing them we found that their nests did not show any significant changes across seasons except that they build deeper chambers during winter. On average the chamber area was 99.17 cc and entrance tunnel were 10.21 cm (Bhattacharyya & Annagiri, 2019). Based on these architectural inputs, meaningful choices were provided to relocating colonies. One nest that was exactly the same as what is found in natural colonies and one that was suboptimal or half the size of what is found in natural conditions. If colonies choose optimal features for their new nest, they should show significant choice towards the former nest. These experiments are ongoing and the colonies choice needs to be quantitatively verified.

**Figure 7.**
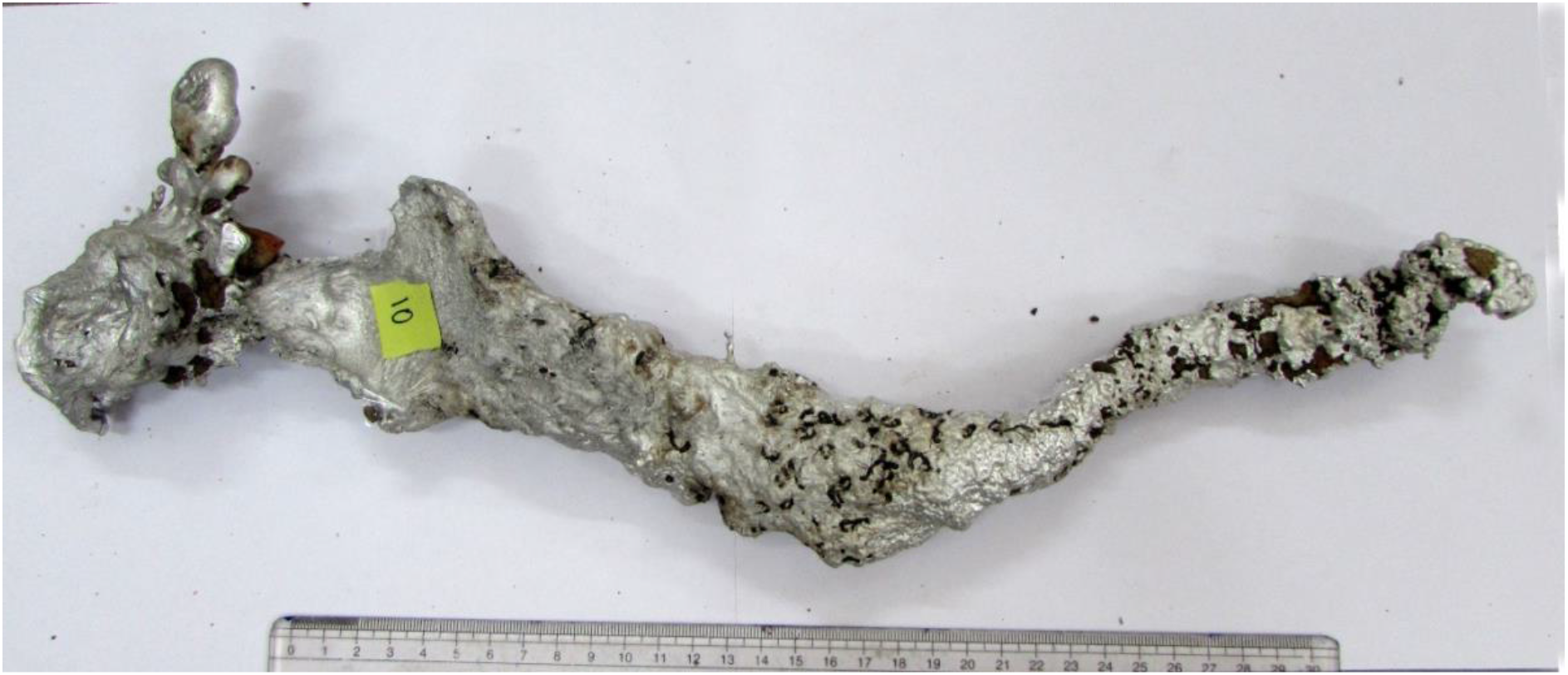
Aluminium cast of *D. indicum*. Entrance is towards left side, followed by the entrance tunnel and then a chamber which has ants encrusted inside it. The secondary tunnel leads deeper into the soil from the chamber. Picture by Kushankur Bhattacharayya,

In addition to the architectural features of the new nest, can the distance over which these ants have to march matter? After all, they have to move hundreds of individuals together with hundreds of immobile brood items; this is going to be complex in terms of logistics and the risk from predators, abiotic stress factors like temperature, humidity and rain can pose. And of course, the neighbours who maybe just waiting to get a chance to take advantage – a topic that will be touched upon in the next section. In nature, colonies relocated to nests which were approximately 1.4 meters from their old nest on average, but they moved into nests that were as close as 30 cm in some cases as well (Kaur et al., 2012). Based on these inputs from the behaviour of these ants in their natural habitat we designed lab experiments to check preferences for different qualities of nests and distances by relocating colonies. Colonies showed a clear preference to move into nests that were already used by conspecifics, they also showed preference towards nests which already had some of their own nestmates as compared to control nests (Kaur & Sumana, 2014). Interestingly when we gave these colonies a trade-off between nest quality and distance, they showed significant choice for short distance, poor quality nest as compared to a good quality nest that was six times further away (Kolay, 2016). Thus, the distance over which they have to travel is important and they minimize this even at the cost of nest quality. This can be because the risk of colony exposure is far greater than time and energy required to restructure their nest structure by digging or accumulating soil and hence, they occupy the near suboptimal nest when they are given a choice.

### Lugging the brood

Unlike other social insects like wasps and honeybees, ants carry their developing young or brood with them when they relocate their nest. This would give them a great advantage as they do not abandon their investments into the next generation, but increase their fitness by bringing the brood with them to the new nest. However, what does it mean to carry hundreds of items in their mandibles while they march? Does this add extra time to the transportation process? To our surprise the answer is no, extra time was not added due to the transport of the brood. When experiments with colonies that did not have any brood and those with brood were compared as they marched over similar distances, under similar abiotic conditions they took similar time. It indicates that this species has incorporated brood transport into the process in a seamless manner (Ghosh & Annagiri, 2022). Further exploration showed that on average 75.3% of all the brood was transported in a coupled manner together with an adult. In other words, followers carry a brood item like eggs, larva or pupa when they tandem run with a leader to the new nest. This couple adult-brood transport was slower but more efficient as it would save an additional trip for the leader as explained in the previous section (Anoop et al., 2021, Kaur et al., 2017). These findings bring up the possibility that the upper limit of the number of brood within a colony can be dependent on the capacity of the adults to transport them during relocation. This is certainly a hypothesis that can be tested. When we increased the brood within the colony, we found that several colonies did not relocate even though they had information about the availability of a new nest (Ghosh & Annagiri, 2022).

### Navigating to the new nest

Have you seen a series of ants walking in a line? Of course, this is a common sight. Several species of ants form a chemical trail along the ground as they perform foraging (Beckers et al., 1989). But all ants do not make chemical trails. Scavengers like *D. indicum* that are solitary foragers and several other ants do not lay a chemical trail as they return back to their nest (Möglich, 1979, Hölldobler and Wilson 1990) each forager has to figure the route back to the nest on her own. While navigation in general is fascinating, having to navigate without the direct aid of nestmates is even more fascinating. Tandem running leaders would have to figure out how to take hundreds of their nestmates from the old to the new nest across heterogeneous terrain. It would be ideal if they can take the shortest route as this would save them both energy and time, but the question is are they capable of figuring out the shortest route? On performing a series of documentation in the natural habitat and experiments in the lab we could address this question. We characterized the path efficiency and speed of these ants. Path efficiency is an index that is derived by dividing the total distance travelled by the ant with its displacement. In other words, if leaders take the straight-line path or the shortest possible path to the new nest, they would score 100 on their path efficiency, if they took a very long path but ended up exactly where they started, they would score 0. On measuring 987 ant’s path efficiency from 8 colonies, in their natural habitat but in a novel surroundings we found that they were remarkably good scoring 83.95 ± 12.45% on average, indicating that they take relatively shorter paths as they tandem run to the new nest (Anoop et al., 2021). What about other ant species, do they also score so high in terms of their speed and path efficiency? Little information is available on path efficiency and speed in the context of relocation in the natural habitat in other species of ants, however some information is available regarding these parameters in the context of foraging in the natural habitat. In *Lasius niger*, path efficiency was reported as 75.38% (Bles et al., 2018) and for Argentine ants that were already familiar with their surroundings it was recorded to be 93.3% (Reid et al., 2011). Next, we wanted to compare the speeds, but the issue is that different species can have different body sizes and thus distance travelled per unit time cannot be directly compared as bigger ants would of course travel more. Thus, body size has to be normalized and this is best done by calculating body lengths travelled per unit time. The walking speeds of *Camponotus consobrinus* is 7.0 bl/sec (Schultheiss et al., 2015), 3.36 bl/sec for *Temnothorax albipenis* (Pratt et al., 2002), 1.41 bl/sec for *Pachycondyla harpax* (Grüter et al., 2018), desert ant like *Cataglyphis fortis* it ranges between 1.55 bl/sec to 213.44 bl/sec (Wahl et al., 2015) and 6.6 bl/sec for *Diacamma indicum* (Kaur et al., 2017). The tandem running speed of *Camponotus consobrinus* is 3.5bl/sec (Schultheiss et al., 2015), *Temnothorax albipenis* is 0.60bl/sec (Pratt et al., 2002), and *Pachycondyla harpax* is 0.84 bl/sec (Grüter et al., 2018). So, in comparison to other tandem running ants, *D. indicum* is the fastest recorded at 4.35 bl/sec on measuring 243 tandem runs in the natural habitat (Anoop et al., 2021).

In the next step we asked a more focused question, can *D. indicum* choose the short path when they are given a choice between a short and a long path? The answer turned out to be yes, majority of the tandem leaders performed almost all their tandem runs on the short path even though they had knowledge of both paths. What happens if things are made a little more complex? Instead of a one decision experiment, if we get the ants to choose at two decision points, would take the short path at both of these decision points? An elegant experiment was performed with trail making Argentine ants by Goss et al., 1989 to answer this question. They found that the inherit volatility of the trail pheromones is sufficient to enable ants to use the shortest path while foraging. But what about ants that do not use trail pheromones and what about ants that need to quickly relocate? Can individual ants with a neural system that is really small figure out the shortest path? Using the same setup as Goss et al 1989, we found that the answer is yes, *D. indicum* tandem leaders carried out the majority of their tandem runs on the shortest path even when faced with two decision points (Figure 8) (Mukhopadhyay et al., 2019). These findings allowed us to conclude that *D. indicum* is capable of minimizing their path while tandem running to their new nest even without the aid of chemical trails (Figure 8B).

**Figure 8.**
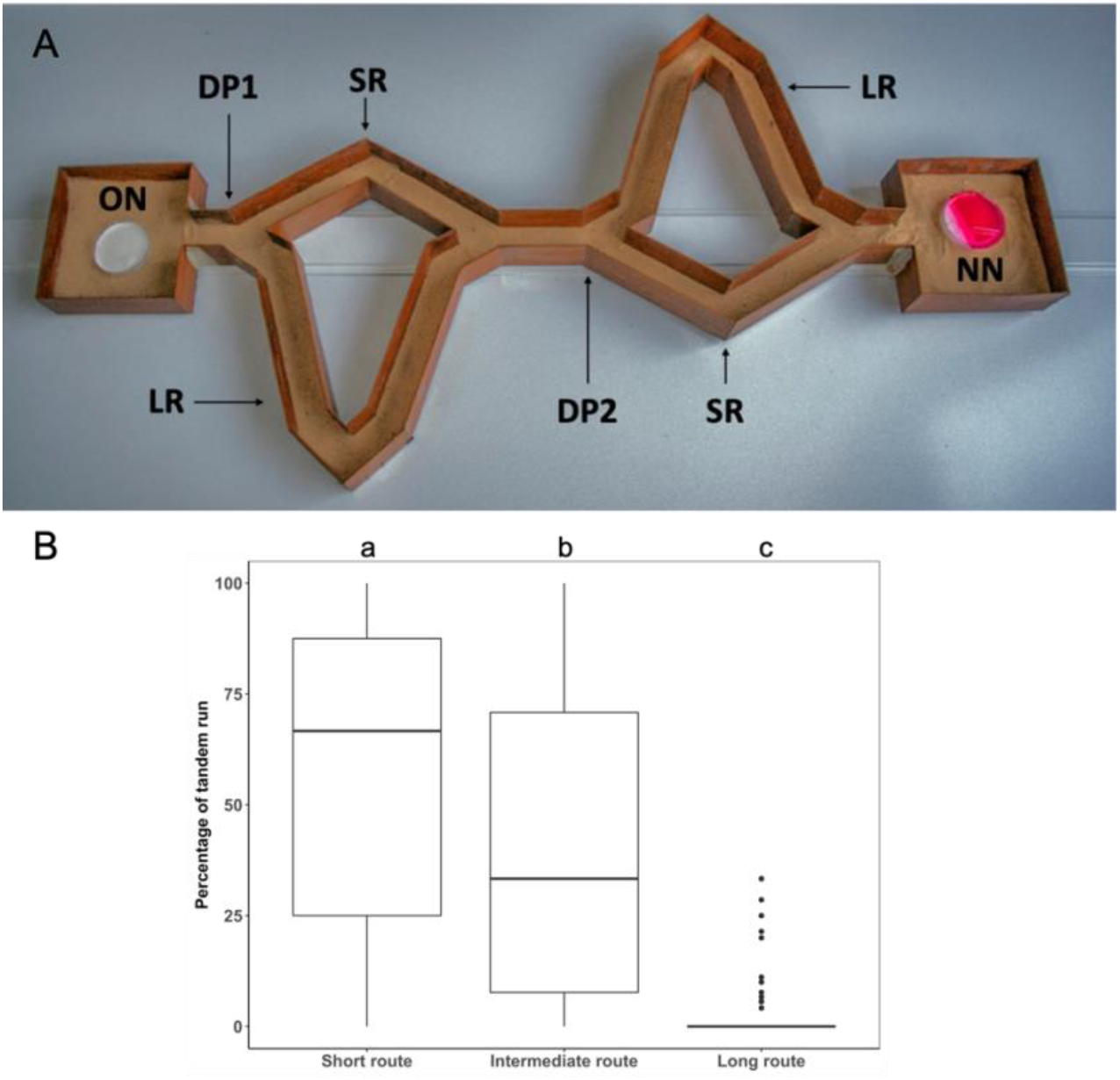
A: The two-decision point maze: This picture shows combination maze different paths and two decision points (labelled as DP1 and DP2) at which ants had to choose. Nests are kept at two ends of the set up indicated as old nest (ON) where the roof is removed in order to induce colonies to relocate and new nest (NN) that is available for colonies to relocate into. In this maze colonies can take the short path (labelled SR) or the long path (labelled LR) at each of the decision points. Thus, tandem leaders can take a combination of these paths during relocation based on their preference for routes, without facing any right or left bias. B) Path choice when a combination of paths was provided. The path choice of individual leaders (N = 77) is shown as the percentage tandem runs that occurred on the short route (SR) at both decision points as short route, the average of the percentage of ants that tandem run on a combination of short route and long route in no particular order as intermediate route and those that tandem run on the long route at both decision points as long route on the X axis. The bold black line inside the box denotes the median value and the box represents the interquartile range.

### Stealing neighbours

Several vertebrates and invertebrates are known to steal. Food is the most common item to be stolen (Breed et al., 2012; Iyengar, 2008). To our surprise we found stealing in *D. indicum*. It was surprising because, tropical ants were not known to steal. Only a few species of temperate ants are known to steal (Hölldobler & Wilson, 1990). Interspecific stealing for slaves has been documented in some ants including *Polyergus* species, in which colonies of one species raid other species to bring back young workers and pupa. These workers then undertake all the day to day duties of foraging, cleaning and raising unrelated ants belonging to the raider’s colony and thus become slaves (Topoff, 1990).

Colonies that become more exposed to both abiotic and biotic components of their habitat were seen to be particularly vulnerable to theft in *D. indicum*. In the natural habitat, we found that 37.1 % of the brood is stolen from relocating colonies by neighbouring conspecific foragers. This quantification was performed by making colonies with marked pupa relocate in the natural habitat in which conspecific colonies resided nearby (average distance 1 meter) and recollecting these colonies after a couple of days to count the number of pupae remaining in the colony. The missing pupa could either eclose into adults or they were stolen. Newly eclosing adults can be identified as they have brownish legs; thus, they can be accounted for upon recollection of the colony. These numbers were contrasted with control experiments in which similarly marked colonies were made to relocate in the natural habitat in which conspecific colonies were not residing in the vicinity and they couldn’t access the focal colony effectively avoiding stealing. We found that almost all the stolen items were pupa the last development stage in the lifecycle of ants. Pupa is the most invested brood item within the colony and is also the most valuable item as it will eclose to produce the next generation of adults either a worker or a reproductive. On following the fate of the stolen pupa in the thief’s colonies across 371 cases in 8 replicates, we found that they are not a food item as none of them were consumed. They were maintained within the brood pile of the thief’s colony and they eclosed to become slaves (Paul et al., 2016). Adults seem to have a strong preference to collect pupa whenever they come across the same and bring it back to their nest. On placing a petri plate with different brood items in the natural habitat of these ants we found that within 30 minutes neighbouring *D. indicum* foragers collect more than 95% of the pupa available with the same individuals doing multiple trips. In lab experiments we delineated the process by which thieves steal and found that collecting unattended pupa within the host’s nest, being undetected and a speedy exit are key factors that contribute to the thieves’ success (Paul & Annagiri, 2019) and thus constituted the tricks of the trade.

**Figure 9:**
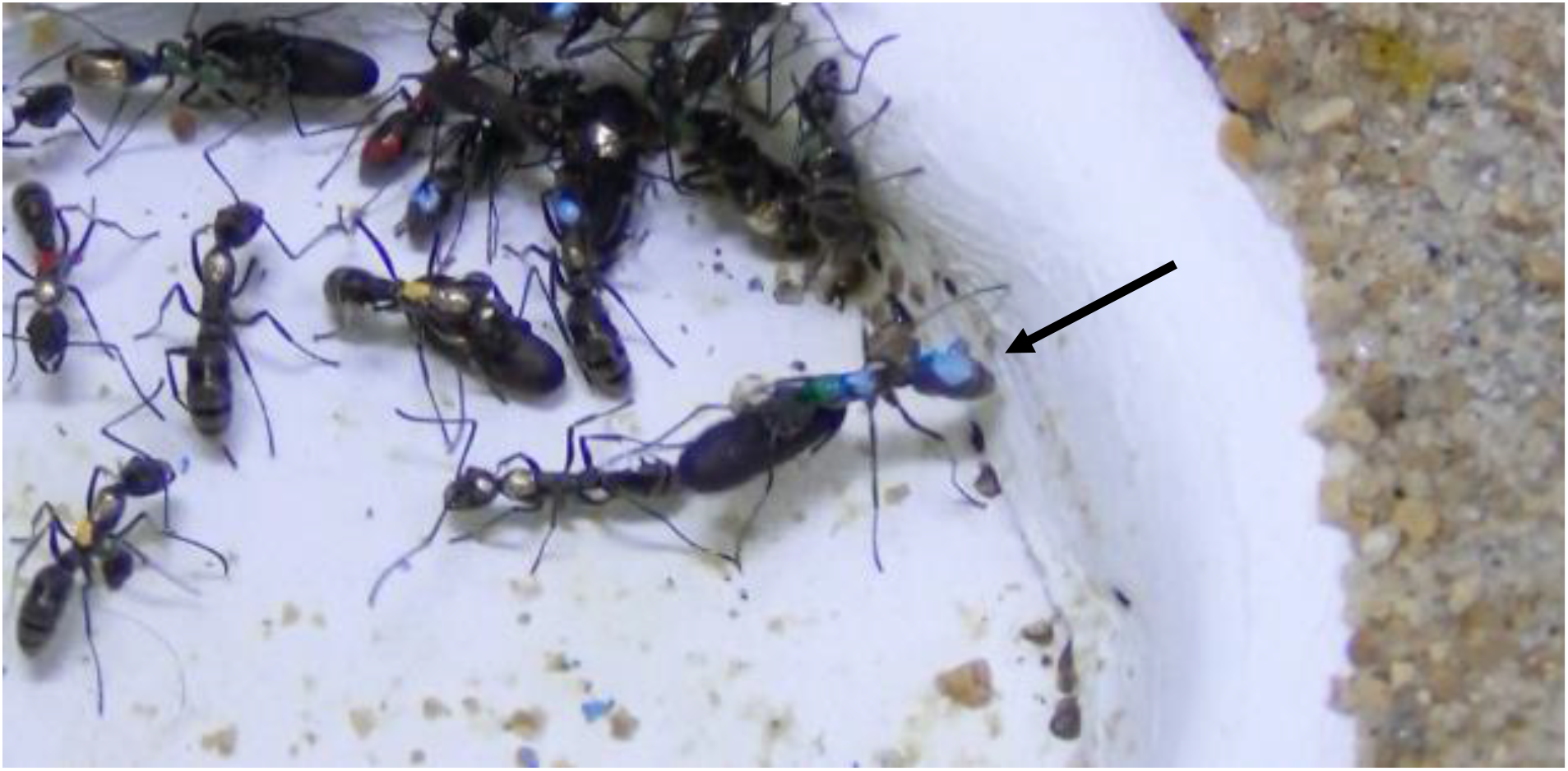
Stealing caught on camera - The ant marked as Green blue blue is stealing a pupa (indicated with a black arrow) from another colony while they are trying to relocate into a new nest.

Currently, it is believed that stealing operates to not only supplement food requirements but it also helps in increasing the workforce and chance of survival for colonies, as seasonality based constraints impose a strong selection force in the temperate regions. (Pollock & Rissing, 1989). However, with these findings we hypothesize that stealing has evolved earlier and perhaps provides distinct advantages even when seasonality is not a major constraining factor in the tropics. Further, given that foragers have a strong preference to collect pupa as highlighted by one individual who was seen to make 18 trips within a single relocation to steal pupa, this affinity for pupa can be used as an incentive to perform many other kinds of experiments. Experiments in which pupa can be used as a reward or unconditional stimulus for conducting learning and memory related studies.

## Branching out

Even though I started off this particular journey by trying to understand what a pair of ants were doing it has become clear that I serendipitously got to open the proverbial pandoras box with this question. Marching along with these ants to their new nest we have had the opportunity to discover several new things by designing and performing really simple experiments that only require a camera and a basic computer in terms of equipment and lots of perseverance, hard work along with it. Coming up with a clear hypothesis and control experiments, workable protocols has been both fun and exhilarating. Doing these experiments with my amazing students has been some of the best times, but publishing them is another story!

In this article we wanted to give a perspective of how ants have evolved to solve basic challenges threatening their colony’s survival and fitness during relocation. This is essential to understanding the lives of these superorganisms. Through the last decade we have started understanding relocation dynamics for this species, but remember there are several thousand species of ants within our country itself and about 14,000 species across the world. To date researchers understand the goal-oriented task of relocation in only a handful species in any detail. Even within *D. indicum* we have exclusively worked with the population found in Bengal, but this species occurs in many other parts of our country and Sri Lanka and possibly Japan as well. What about these populations will they mirror these same dynamics or has the local biotic and abiotic factors changed certain aspects of the relocation dynamics? In addition to understanding relocation dynamics *per se* our lab has now graduated into using relocation as a platform to understand different aspects of the life of these ants. To list a few, the intra and inter species interactions regarding dwellings and the effects of monsoons on the natural history of these organisms. As you can imagine, answering one question only gives rise to at least another five questions while conducting research. We are currently pursuing the answers for some of the following questions. How do they deal with traffic jams? Do they rescue buried nestmates and brood, if yes, what is the precision with which they conduct this rescue? Do they show any preference during rescue? Do they store information about the location of their new nest in their memory? How long does this memory last? When is this memory formed?

This is by no means a complete listing of our findings nor a thorough review of what is already known in the field. Instead, this review intends only to give a little glimpse of the research we have done so far, written in a manner that can be easily understood. In the long run, we hope to understand this ant in particular and societies in general along our journey.

## Declaration

On behalf of all authors, the corresponding author states that there is no conflict of interest. We have conducted all our experiments in accordance with the guidelines that are applicable to working with the model organism in our country. We have collected colonies from areas close to human habitation that is not part of any protected areas or forests. Colonies were maintained in the laboratory with ad libitum food and water and after the experiment they were released back in the habitat from which they were collected.

## Acknowledgements

We thank Purbayan Ghosh for entering information from the incoming colony register into excel. We thank Basudev Ghosh for his help in maintaining colonies in the lab and collecting them. We thank Subhashis Halder and Kushankur Bhattacharyya for their excellent photographs. We thank Sudhi Rao for constant supply of Testors paints and feather forceps. We thank Indian Institute of Science Education and Research Kolkata for the support they have given and SERB for the two grants SR/FT/LS-179/2009 and EMR/2017/00147 over the years. We also thank University grants Commission, INSPIRE and CSIR for giving fellowship to the Ph.D. students who have worked in the lab over the years. We also thank two anonymous reviewers for their suggestions which has helped us improve our manuscript.

